# Analytic Model of fMRI BOLD Signals for Separable Metrics of Neural and Metabolic Activity

**DOI:** 10.1101/573006

**Authors:** Charles D. Schaper

## Abstract

The applications of fMRI (functional magnetic resonance imaging) are broad covering diagnostic and clinical extents of brain function, which involves the analysis of BOLD (blood oxygen level-dependent) contrast signals. The BOLD signals are sourced from both neural and metabolic functions, and thus to enable a detailed examination of fMRI studies, methods are sought to separate the neural and metabolic functions, such that the neural component, which is often the metric of interest, can be independently examined, especially in relation to neural connectivity. In this work, a modeling approach is developed that separates the neural and metabolic functions from the overall BOLD signal. The newly developed model is initially developed within a linear framework and demonstrates excellent comparison in data fit at 97.4% to the three Gamma function, which has been widely used to characterize fMRI BOLD experimental data. The neural component of the model formulation is comprised of a proper transfer function of two poles and two zeros, and characterizes the salient features of the BOLD signal, including the initial dip, peak, undershoot, and stabilization period. The linear model is extended to characterize nonlinear fMRI BOLD signal responses through the integration of saturation functions to both the leading and trailing edges of the signal. The nonlinear model representation not only explains the muted response in amplitude and oscillations, but also explains nuanced oscillations during the hold and settling phases of fMRI BOLD responses as exemplified in comparison to published data of sensorimotor responses. Further, the newly developed decomposition is derived within a framework for modeling neurovascular coupling environments, and thus lends credibility to the modeling framework. In developing the decomposition of the neural and metabolic transfer functions, it is a conclusion that the BOLD signal correlates very well with the fast dynamics associated with neural response to external stimuli.

**Graphical Abstract:** 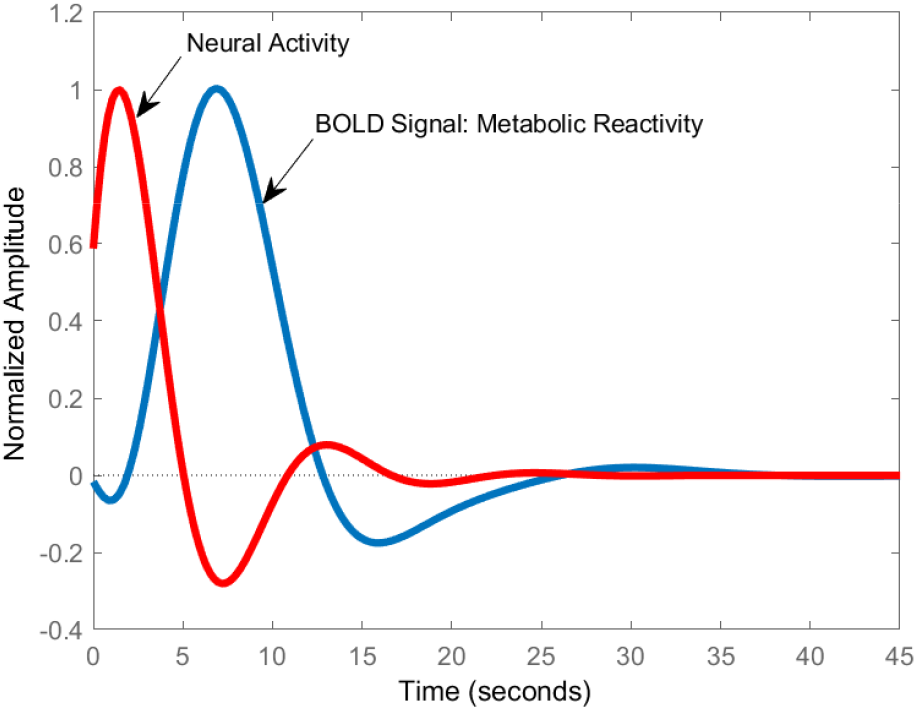

The normalized impulse response of the BOLD signal and the corresponding neural activity according to the newly developed model. There is a correspondence of the critical points for the oscillatory response of the neural function and metabolic reactivity, including the initial dip, peak and subsequent undershoot. Thus, the BOLD signal is a correlated representation of the underlying neural response.

## 1 Introduction

The applications for functional Magnetic Response Imaging (fMRI) in relation to brain function are numerous, profound, and range widely including studies on dementia [28], drug dependency [16, 5], brain-computer interface [31, 13], ADHD clinical testing [14], psychiatric disorders [12], stress [33], neurological disorders [8], parkinson’s disease [27], aging [24], and cancer[26].

As fMRI produces a BOLD (blood oxygen level-dependent) contrast image signal, both neural activity and metabolic reactivity are incorporated within the signal, which is thereby influenced by blood flow [1, 21]. To enhance the signal to noise ratio of the signal, and to aid in its understanding, modeling techniques have been developed, although the exact cellular processes comprising the neurochemical contribution of the BOLD signal in mapping neural activity in fMRI responses to external and internal stimuli have yet to be exactly determined [15, 18].

It is an objective of this work to separate the neural activity and metabolic reactivity from the BOLD signal, and subsequently develop a decoupled analysis of the neural and metabolic components to improve the accuracy of the correlations, as well as a detailed examination of the underlying neural and metabolic mechanisms driving physiological function. Model-based techniques for evaluating neural contributions of the BOLD signal include[17, 11, 6]. Neural coupling to cerebral blood flow and oxygen changes in cerebral metabolic rate have been developed but there is no consensus on the exact neural activity, and nature of the neural forcing function, that drives the hemodynamic response [2]. To further the investigation of the neurovascular coupling, contrast agents [4] have been deployed to evaluate the impulse response of neural activity and recent modeling approaches have been developed as well [9]. Further, in promoting a basic understanding of the neural response, it is surmised that activity at the synaptic cleft plays a role in generating BOLD signals as examined in Alzheimer’s disease and Ketamine responses for example [23, 22].

Furthermore, in relation to the challenges of characterizing fMRI BOLD signal as neural and metabolic components, a challenge is that under nominal external stimuli driven responses, the brain function can transition from a linear to a nonlinear response, in which the measured signal is not a linear superposition of impulse functions, but rather is attenuated in amplitude and transient dynamics. Consequently, modeling techniques of BOLD signals have been extended to include such nonlinear features [19, 32, 7], and deployed for example to analyze fMRI data of the visual cortex [20] and for dynamic modeling of applications in connectivity [25].

In this work, to extract the neural and metabolic activity as dynamic metrics from the fMRI BOLD signal responses, a recent framework for neurovascular coupling of brain function [29] is utilized, which enables a transfer function characterization of the BOLD contrast image signal ofr linear responses to internal and external stimuli. This transfer function description is then separated into neural and metabolic functions and analyzed to show excellent fits in comparison to existing model descriptions of the response, such as three Gamma function model. After developing the linear relations, extensions are then developed to characterize nonlinear responses by examining saturation at the neural level. It is compared against experimental data and models, including a modification of the balloon model, from a prior study of Glover[10].

## 2 Methods

To construct metrics characterizing both the neural and metabolic components of fMRI BOLD signals, a continuous time model is constructed with reference driving inputs as external stimuli as well as from neighboring neural inputs. To start the analysis, a linear framework is used. To expand the range of BOLD signal responses, a nonlinear model is incorporated in which response functions associated with the synaptic cleft are incorporated, effectively limiting the speed and amplitude of the system response.

### 2.1 Schematic

As a representation of the proposed model developed for characterizing fMRI signals, a schematic is shown in Figure 1 in which the relation between the incoming signals, which for this case are derived from exogeneous or event-driven stimuli as well as from the axon terminals of neighboring neurons. To fuel the transmission of incoming neurotransmitter signals, as well as the overall neural response, metabolic chemicals are modeled and described within the overall system response. For purposes of analysis, the metabolic signals are considered as a mediator in conducting the neural response and are also modeled in direct proportion to the BOLD signal response. Neurovascular coupling from external stimuli is comprised of signals originating from neurotransmitters that interacts with a metabolic components in generating an input signal to the axon. Subsequent interaction with metabolic materials then result in an output from the axon terminals.

**Figure 1:**
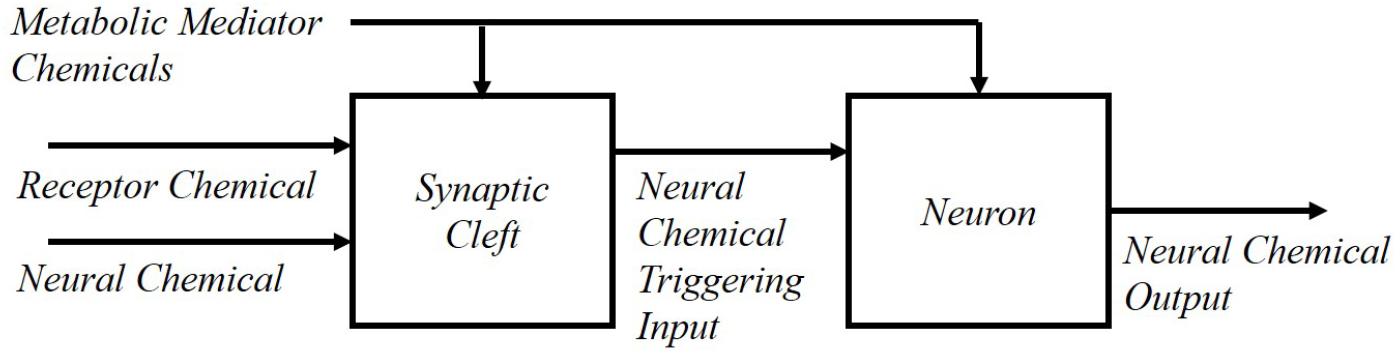
To develop a separable model of the neural and metabolic functions comprising the fMRI BOLD signals, a schematic of the response is utilized in which receptor and neural chemicals interact at the synaptic cleft under metabolic interaction with a mediator chemical species, which is considered proportional to the BOLD signal. Subsequently, a signal via the membrane and in association with metabolic mediator chemical species produces the input charge to induce a neural signal.

### 2.2 Control Law

A control law is developed that describes the trigger forcing a neural action in response to chemical activity in the synaptic cleft as presented in Figure 2. The input and output variables at the synaptic cleft are introduced as *Q*(*t*), which signifies the metabolic mediator chemical that is modeled as proportional to the fMRI BOLD signal; the input from neighboring neurons is denoted as *Y*(*t*); and the input from exogenous signals is denoted as *Y*(*t*). After interaction within the synaptic cleft and at the post-synaptic membrane substrate, a signal *U*(*t*) is generated as input to the neuron. It is noted that this expression and control policy is in-line with that of [29], which was developed to characterize neurovascular coupling in explaining feedback control of body temperature in response to a chemical mediator and neural signals, especially in response to infectious agents.

**Figure 2:**
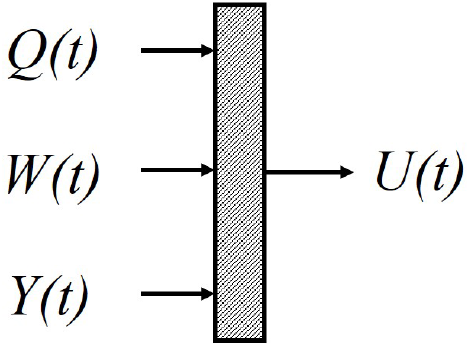
The input and output variables at the synaptic cleft are introduced as Q(t), which signifies the metabolic mediator chemical, which is proportional to the fMRI BOLD signal, the input from neighboring neurons Y(t), and the input from an exogenous signal Y(t). After interaction within the synaptic cleft and interacting on the post-synaptic membrane substrate, a signal U(t) is generated as input to the neuron.

The control law incorporates the rate of change as an input variable because the process of membrane activation involves a diffusion process, which is governed by spatial concentration gradients. Thus, the time derivative of the chemical input species relates to the adjustment of the concentration gradient driving the membrane charge. Thus, the control law is considered to be of the general form as described in Equation (1)

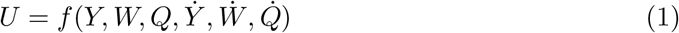

where *U* is an effective signal into the neuron at the post-synaptic membrane associated with the generation of action potentials, *Y* denotes the chemical neurotransmitter input from internal sources such as axon terminals, *W* represents the neural signal associated with external stimuli such as neuroreceptors, *Q* is the metabolic chemical species that interacts with *Y* and *W* to trigger responses of the postsynaptic cell and is proportional to the oxygen level as per BOLD imaging, *Ẏ* ≡ *dY*/*dt, Ẇ* ≡ *dW*/*dt* and 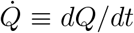, which are the rates of change associated of the neurochemical responses.

Linearizing Equation (1) about a nominal operating point, results in the following representation

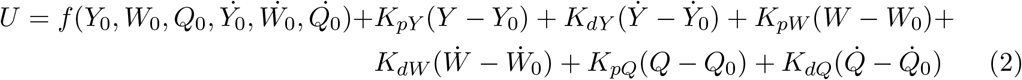

of which the controller gain factors, critical in defining performance, are given by:

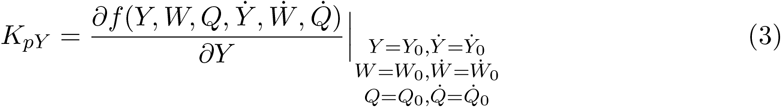

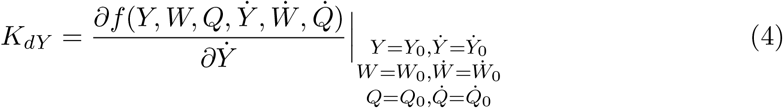

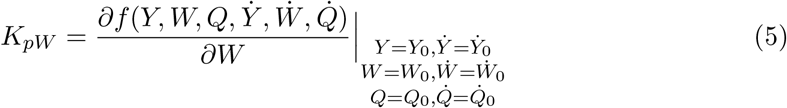

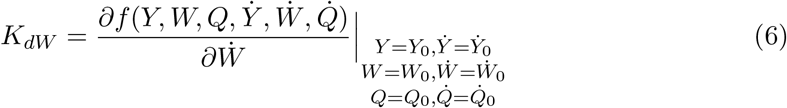

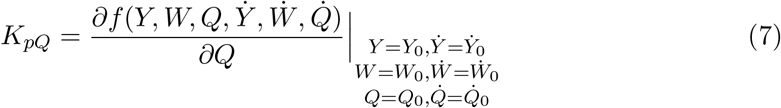

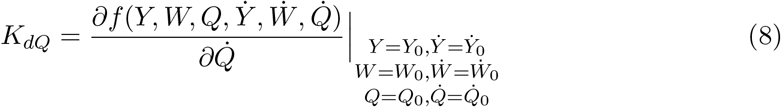

Deviation variables are defined as Δ*U* = *U* − *U*_0_, Δ*Y* = *Y* − *Y*_0_, Δ*W* = *W* − *W*_0_, Δ*Q* = *Q* − *Q*_0_, and thus the control law expressed after substitution into Equation (2) is represented as

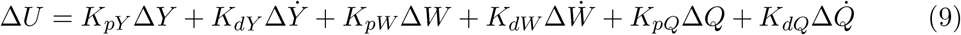

The control law expression is proportional to the summation of signals due to the neighboring axon terminals and external stimuli in which a metabolic chemical is integrated to energize the actions. While the gains were determined in a linearization of the basic form, it is reasonable to consider that the reaction rate is proportional to a bilinear association of *Y* and *Q*, and with *W* and *Q*, which when linearized involves a decoupled linear format.

### 2.3 Neural Response

The neural response *X* to the input signal *U* as mediated by metabolic chemical *Q* is represented by the form

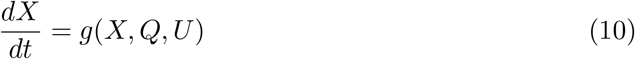

A linear dynamic model is obtained from a Taylor Series expansion of Equation (10) and then neglecting all terms of order two and higher, the expression for the neural response is represented as:

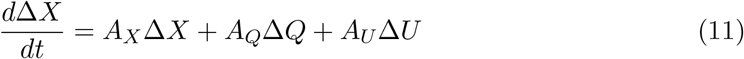

where

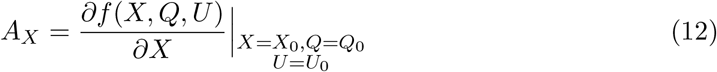

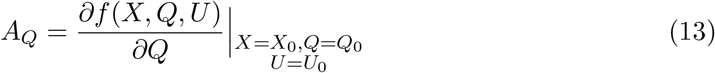

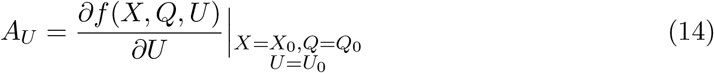

Equation (11) is a linearized model which maps the neural response from inputs generated via the control law and energized by metabolic mediator materials. It is typical of models used for linear time invariant analysis of dynamics.

### 2.4 Neurovascular Coupling

To couple a metabolic function with the neural signal, the dynamics are expressed as follows:

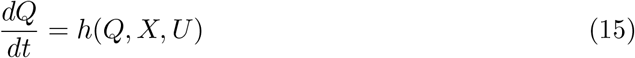

and in developing a linearized representation of Equation (15), a Taylor Series expansion truncated after the first term yields:

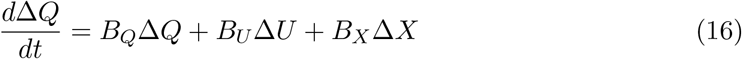

where

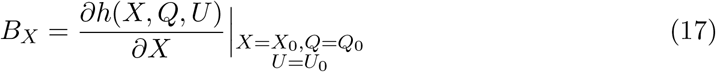

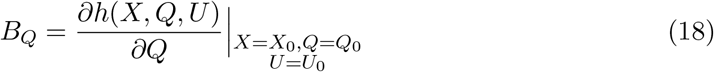

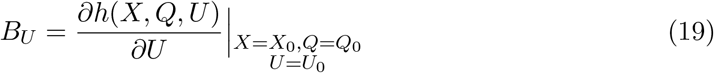

which is similar in form to that of the neural dynamics, Equation (11). It is of interest to note that in this model, the mapping to the neural vascular channels supplying the metabolic chemical are considered as a constant source from which *Q* is sufficiently supplied to produce the necessary metabolic functions to drive the neural response.

### 2.5 Linear Dynamic System Response

In forming a set of linear, time-invariant models to describe the neural and metabolic actions, the Laplace transform of Equations (9), (11), (16), in 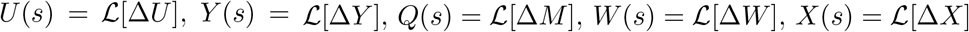,

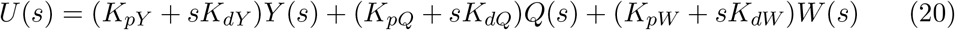

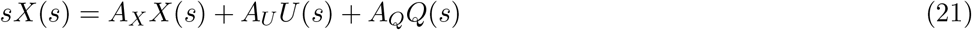

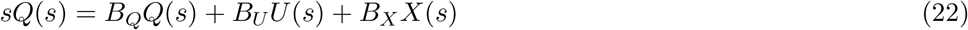

To map the external stimuli signal *W*(*s*) to the BOLD response signal *Q*(*s*) when the neighboring neural signal *Y*(*s*) = 0, the intermediate signal associated with the neuron action potential *X*(*s*) is considered,

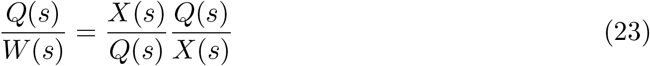

The expressions for the functions from Equations (20), (21), and (22) are,

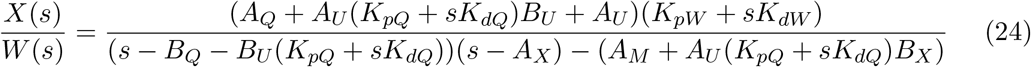

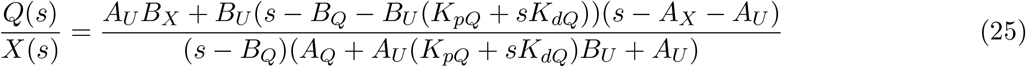

It is noted that the order of the model from *Q*(*s*) to *W*(*s*) is four poles and four zeros. Thus, to determine the contributions of the BOLD response signal from the triggered neural process *X*(*s*) from an external stimuli *W*(*s*) based upon measurements of *Q*(*s*), the strategy is to determine an overall map of the transfer function *Q*(*s*)/*W*(*s*) as a four-pole, four-zero expression. Because the metabolic function is known to contribute more slowly to the BOLD response than that of the neural function, the identified expression is decomposed into two fast poles and two fast zeros combinations associated with the response *X*(*s*)/*W*(*s*) and the remainder of the identification transfer functions is then selected for the response *Q*(*s*)/*X*(*s*). This construction is described in the following representation:

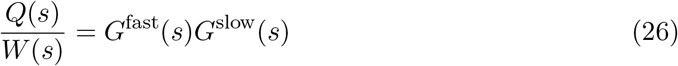

in which *X*(*s*)/*W*(*s*) = *G*^fast^(*s*) and *Q*(*s*)/*X*(*s*) = *G*^slow^(*s*).

### 2.6 Connectivity

For the connection of neighboring neurons, the relation between the internal stimuli of transmitting fibers or of neighboring axon terminals in triggering an electrical nerve impulse is described by the dynamics of the process as,

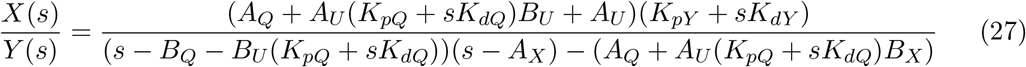

with Equation (25) providing the relation of *Q*(*s*) to *X*(*s*).

### 2.7 Nonlinear Modeling

It is well known that the BOLD signals exhibit nonlinear behavior, especially in cases where the stimuli are presented in rapid fashion, in which a simple additive response of the impulse function does not adquately describe the observed signals. The signals are much less in amplitude than that which would be expected if the response were linear. For example, as will be seen, whereas the amplitude would increase by a factor of five for a linear response, an increase of only by a factor of two for a nonlinear response. In addition, other factors of the BOLD response, such as oscillations of undershoots and overshoots are damped in comparison to what would be expected in a linear response to an increase in external stimuli strength.

In the proposed model, the source of the nonlinearities is postulated as due to the association of the neurotransmitters from the triggering stimuli, either internal or external, with the post-synaptic receptors on the plasma membrane that convert the electrical charge on the inside of the neural cell membrane. As the amplitude of the input signal is increased and sustained, the receptor channels are occupied by the neurotransmitter and the reuptake speed is too slow to process all of the incoming chemicals, and thus it simply reaches a maximum number of sites. Prior to this *saturation* at the onset of the triggering signal, the incoming signals can be processed quickly even though the rate of initiation may be significantly more than that which is moved into and out of the neuroreceptor field when saturated. The relation of Equation (28) is used to characterize this saturation process,

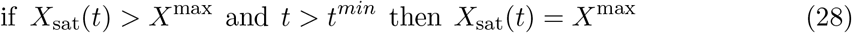

where *X*_sat_(*t*) is the neural response in saturated condition, *X*_max_ is the maximum achievable rate of neural excitation given a saturated state, and *t*^min^ is the time at which the post-synaptic neural membrane first achieves a saturated condition.

For the cumulative effect of neural transmission sourced from axon terminals of the external stimuli and from transmitting neighboring neurons, a certain number of receptor sites will enter into saturation from a linear state; will operate in a saturated state with a build-up of neurotransmitter; or will enter saturation that is subsequently depleted over time. To determine the depletion of the receptor and the neurotransmitter within the synaptic cleft, a model incorporating an initial condition of the neural state *X*(0) and metabolic state *Q*(0) that is gradually depleted is modeled by the expression,

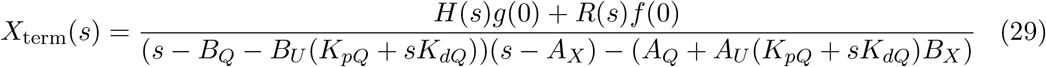

where *X*_term_(*s*) is the neural activity in saturation conditions, and where

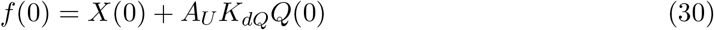

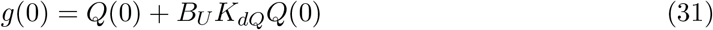

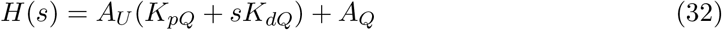

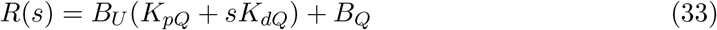

which depends upon the control law parameters associated with *Q*(*s*), that is *K_pQ_* and *K_dQ_*. And as *X*(*s*) is processed, the metabolic function will follow a delayed response, which is considered as a time delay with similar dynamics, that is a first order response, in which the depletion model associated with the metabolic function may be associated as a second order time delay function with one zero as follows:

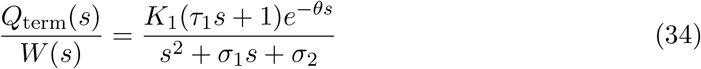

where *Q*_term_ is the portion of the metabolic material associated with the saturated state, *θ* is the time delay, *K*_1_, *τ*_1_, *σ*_1_, *σ*_2_ are determined from the terminating slope as measured by the BOLD signal after the input excitation has ceased and in which the input excitation has been sufficient to saturate the response.

The fractional contribution to the BOLD signal from signals initially unsaturated and saturated can be found according to Equation (35)

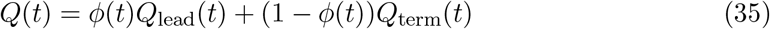

in which the *ϕ*(*t*) weighting function is determined as the fraction of signals in saturated mode and unsaturated *Q_lead_* mode. This parameter is correlated to the membrane diffusion equation of

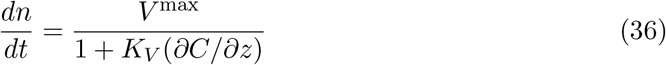

where *dn*/*dt* denotes the rate of change of the receptor, *V*^max^ and *K_V_* relate to the peak and shape of the diffusive function, and *∂C*/*∂z* denotes the concentration gradient. This asymptotic consideration can be approximated as the sigmoidal function

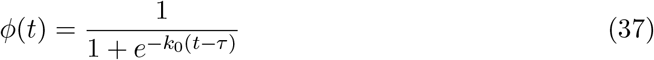

The modeling parameters *k*_0_ and *τ* can be set in accordance with the experimental test configuration.

## 3 Results

To assess the decoupling of the neural and metabolic functions from the BOLD signal, as well as the feasibility of a control law expression that included neurovascular signals, a set of studies were conducted in comparison to models previously fit to experimental data. The rationale is if the newly developed model can accurately track model forms that have themselves been used to accurately fit experimental data, then the newly developed will work at least as equally well. The models selected for this study were the three gamma function, which has been applied to many fMRI studies for linear responses. And to characterize the nonlinear performance, the model selected was that of Glover [10] which characterized a nonlinear response of experimental data, and was compared to the well known Balloon model [3]. In addition, data previously measured in [10] was used directly as well to assess the ability of the newly developed model to adequately filter experimental noise.

### 3.1 Linear Neurophysiological Responses

To examine the linear function response, the three gamma model described in [30] was used with the following parameters, which corresponded to a sensorimotor response.

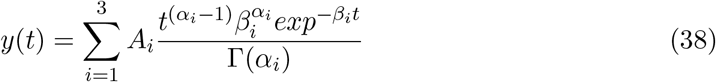

where *A*_1_ = −0.2; *A*_2_ = 10; *A*_3_ = −3.6; *α*_1_ = 1.5; *α*_2_ = 6.6; *α*_3_ = 15; *β*_1_ = 0.8; *β*_2_ = 0.8; *β*_3_ = 1;

An impulse response was generated and the data used to determine a four pole, four zero model using Matlab System Identification toolset. The results are depicted in Figure 3, which shows an excellent fit to the data model of 97.4%. The salient features of the BOLD response are captured very well including the initial dip, overshoot, undershoot, and stabilization period as well as the model gains.

**Figure 3:**
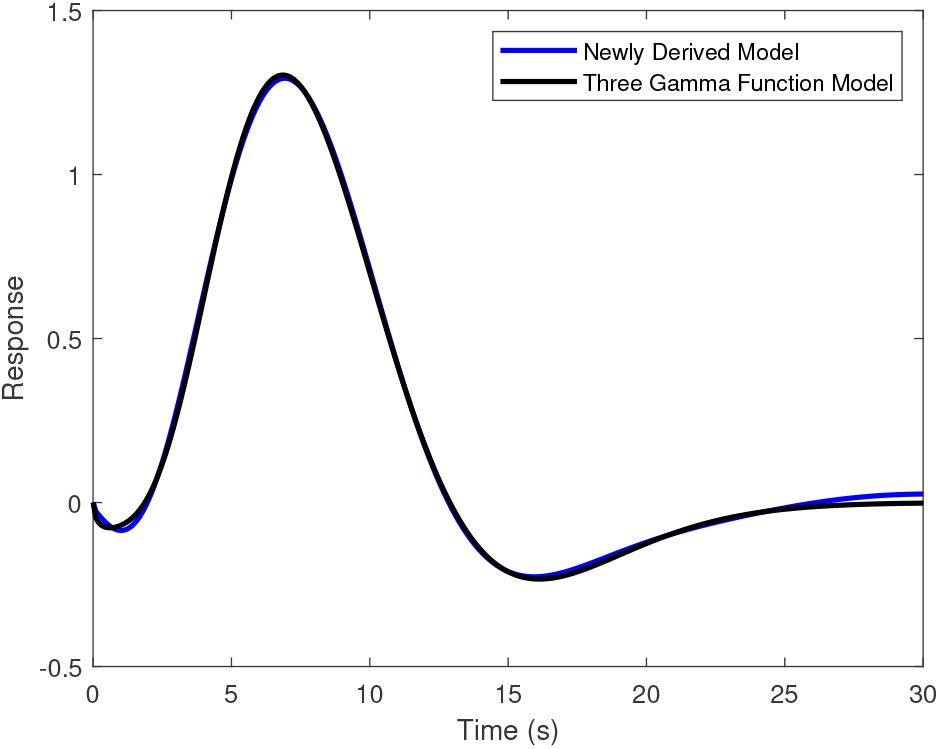
The impulse response of the newly developed model is compared with that of the three gamma function indicating a model fit of 97.4%, which indicates excellent agreement, and includes a match in the initial dip, peak and other features of the oscillatory nature as it settles to baseline.

The utility of the proposed model is shown here by decomposing the result into fast and slow modes, and assigning the fast modes to the neural response as described in Equation (26). The resultant transfer functions are presented in Equation (39)

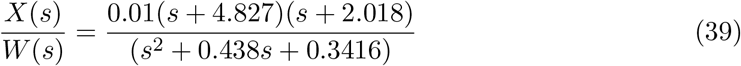

And for the transfer function from the neural component to the metabolic function, the result is thus

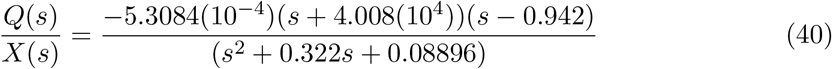

The impulse response of *X*(*s*)/*W*(*s*) is indicated in Figure 4 which shows a rapid initial response at the onset, due to the derivative nature of the model, followed by an increase and then a reduction back to baseline.

**Figure 4:**
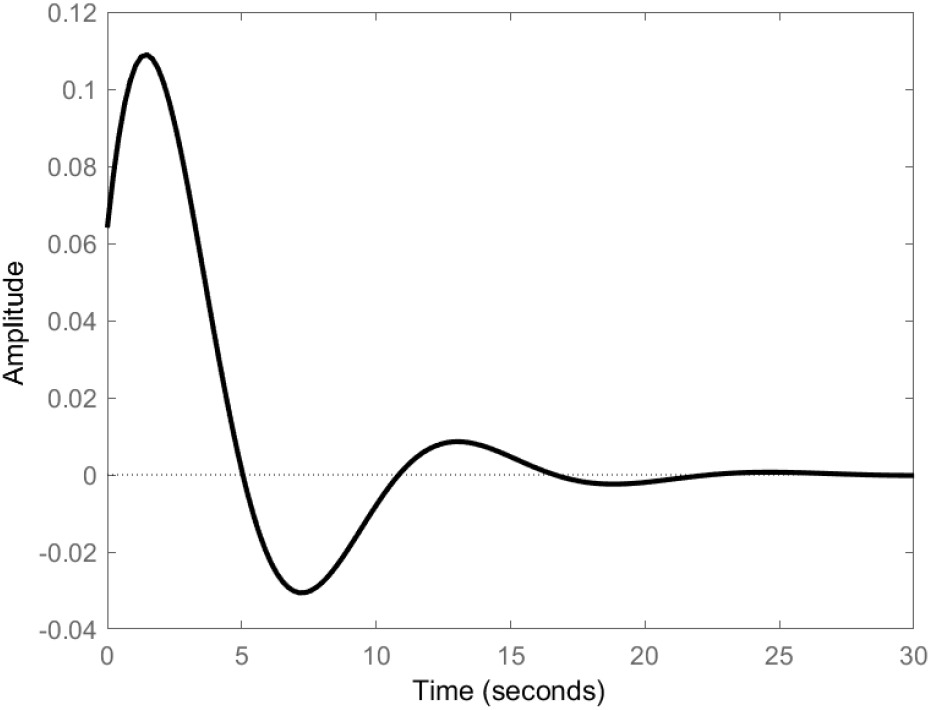
The impulse response of the neural response according to Equation (39). Note the immediate activity of the neural response due to the derivative term of the model, while the oscillatory nature of the BOLD signal is also seen in the neural response, including initial dip, peak, and undershoot.

Comparison of the neural activity and the observable BOLD signal is presented after normalization such that the maximum amplitude for both signals is 1 in Figure 5. The result shows correlation of the neural transitions to the BOLD signal: for example, the initial dip correlates in time to the rise of the neural signal, and the maximum signal of the BOLD corresponds to the completion of the initial cycle of the BOLD signal. Other correlations are seen via oscillations in the tail portion of the response curves. It is noted that the neural signal completes it cycle largely in the first 4 seconds to reach ± 10% of its final value of 0, whereas the BOLD signal takes about 12 seconds.

**Figure 5:**
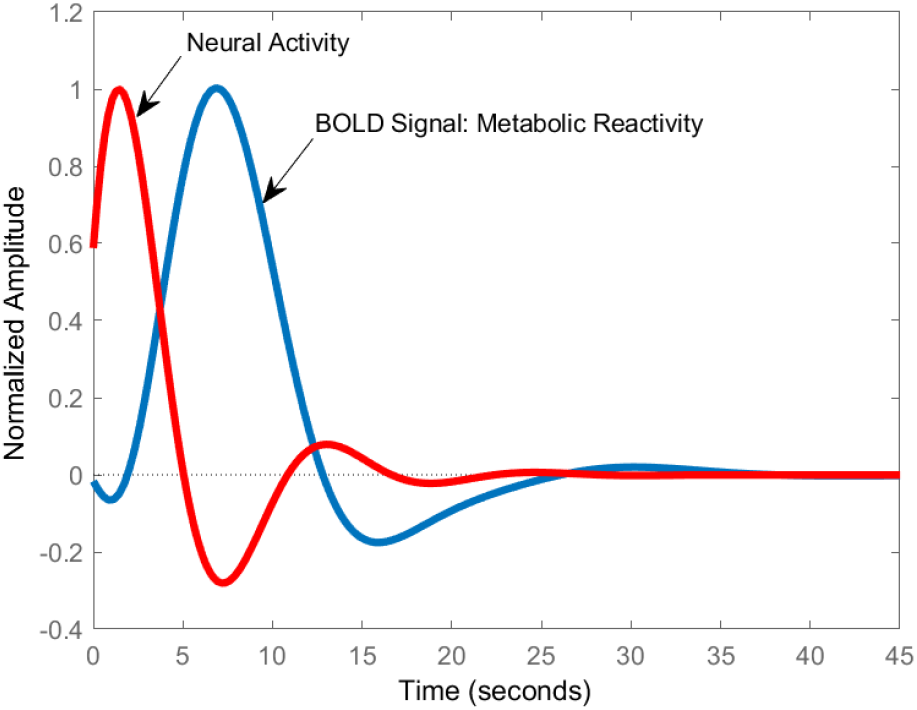
The comparison of the impulse response comparison of the neural component X(s) and the BOLD signal component Q(s), with both signals normalized such that the maximum amplitude is 1. There is correlation in the oscillations of the neural and BOLD signals, including the initial dip, peak and undershoot. The metabolic function of the BOLD signal is an inverse respone relative to the driving neural function. The impulse response for neural activity precedes the BOLD signal by a factor of approximately 3 and is initiated instantaneously via the external stimulus.

The rationale for the impulse response relative to the observed BOLD signal is that there is an initial rapid response of the neural component at time 0 when the input driving signal is initiated. Rather than immediately settle after the driving function is terminated however, there are substantial dynamics taking place afterwards. It is seen that the neural signal rises possibly as more neurons within the vicinity of neurotransmitter release potentially due to diffusion processes are recruited into processing the neurotransmitter load. There is an initial dip in oxygen due to depletion because of the timing at which the diffusive nature of oxygen can be transported to the sites of activity. The neural response then subsides as there is reuptake of the neurotransmitters, and an undershoot due to the depletion of the oxygen supply relative to its value at resting state which makes the neuron less active than that at its resting state. The neuron and oxygen levels are finally reestablished at steady-state. It is noted that the time scales involved in this process are surprisingly long relative to the very short duration of the action potential, which is on the order of 10 to 100 milliseconds. In the comparison of the shape and critical points of the proximate respone curves, it is a conclusion that the BOLD signal is an exceptional correlator to the neural response.

The frequency plots of amplitude and phase of the transfer functions of *X*(*s*)/*W*(*s*) and *Q*(*s*)/*W*(*s*) are presented in Figure 6, which indicates a sharp resonant frequency and rapid roll-off associated with the neural response and a slow roll-off shown for the metabolic response. The metabolic response is seen to be larger than the neural signal in basic response, and that is due to the allocation of gain function from the overall response transfer function. The selection was taken as 0.01 because larger values result in instabilities when proceeding from neural to neural connection, as presented shortly. Lower values in gain can be selected for the neural activity without issues of stability arising.

**Figure 6:**
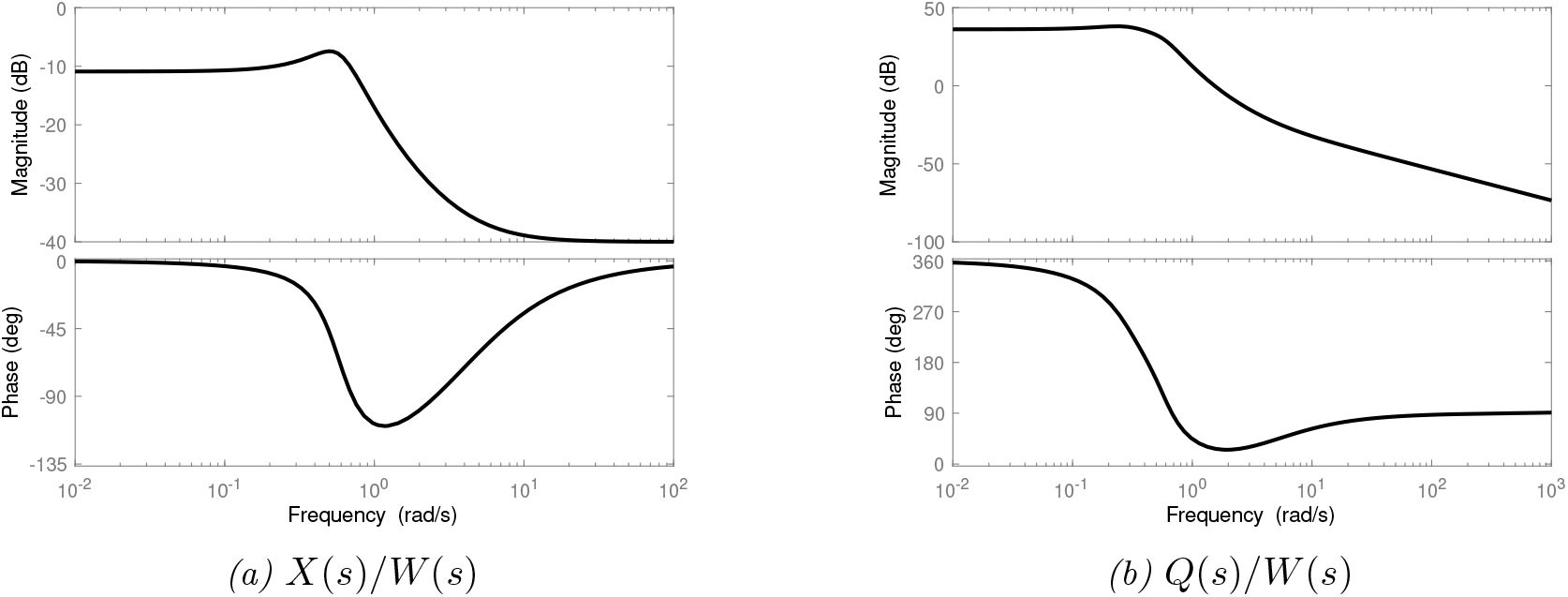
The frequency responses of the neural and metabolic transfer functions. The result indicates a resonance frequency in the neural response of approximately 0.5 rad/sec and a sharp roll-off at higher frequency inputs, especially in relation to the metabolic function, which in an electrical system would aid in preventing cross-talk between adjacent channels.

To evaluate connectivity, the model of Equation (39) is processed multiple times to determine the progression of the neural signal if there were no resistance between the connections, such as potentially in a gap junction. The result is indicated in Figure 7a, which shows a quick reduction in signal strength from the initial impulse. The strength of the progression defining neural connectivity is dependent upon the gain selected for the impulse signals, which shows a four-fold reduction in amplitude from connection to connection, and a filtered response of the second impulse in comparison to the initial response. To investigate a change in gain to achieve an increase in neural connections, the result is shown in Figure 8b where the gain is increased by a factor of three from that of Figure 7a. The result is dramatic as there are more powerful impulses in the second, third and fourth connections; however there is an instability in the response such that by the tenth connection, there is significant increase in the response. This is caused likely by the resonant frequency stimulation which is increasing faster than the other modes of the response can be reduced.

**Figure 7:**
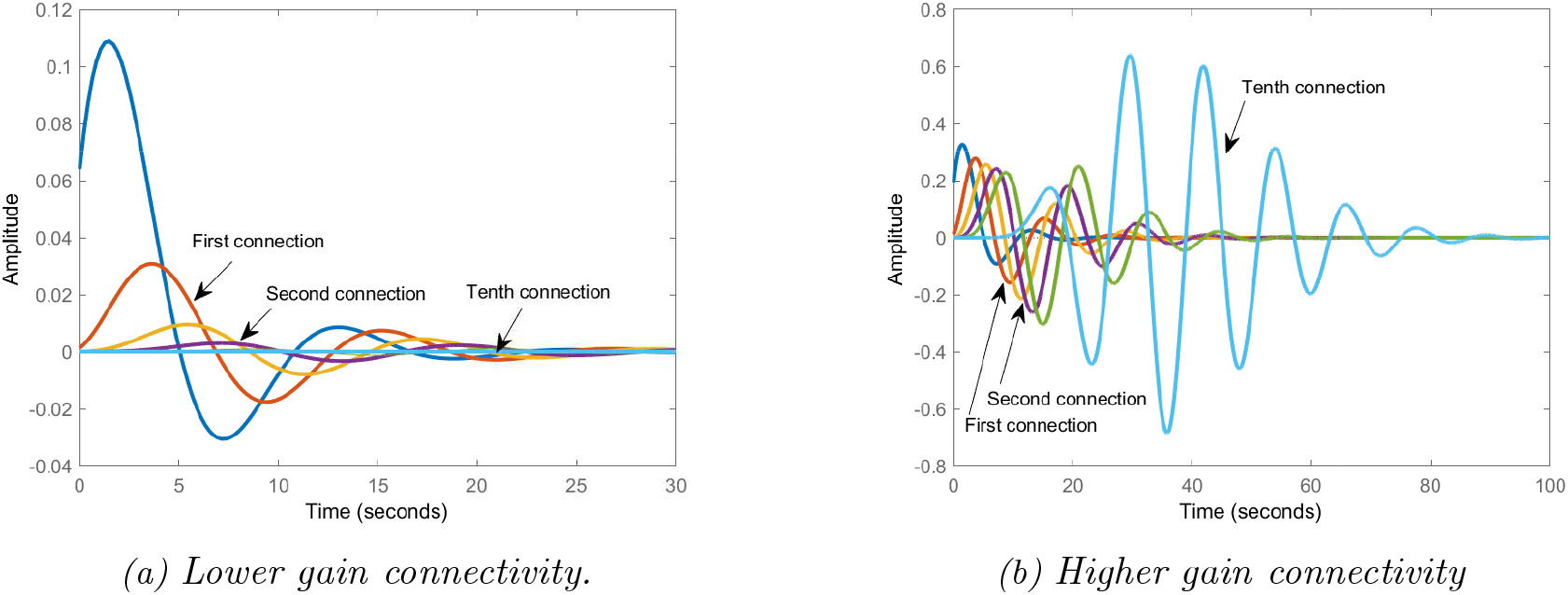
Investigating the connectivity of neuron to neuron, the gain parameter, which is a parameter in the models, was adjusted to determine the nature of connecting a neuron to neuron, without delay or attenuation from one neuron to the next. Due to resonance, it was determined that a low gain was required, in order to avoid the generation of an unstable response due to the resonant frequency. It was determined that less than ten sequential connections for this function would be the limit as a higher gain from neuron to neuron, which would aid in the number of neurons connected sequential, would result in instability.

**Figure 8:**
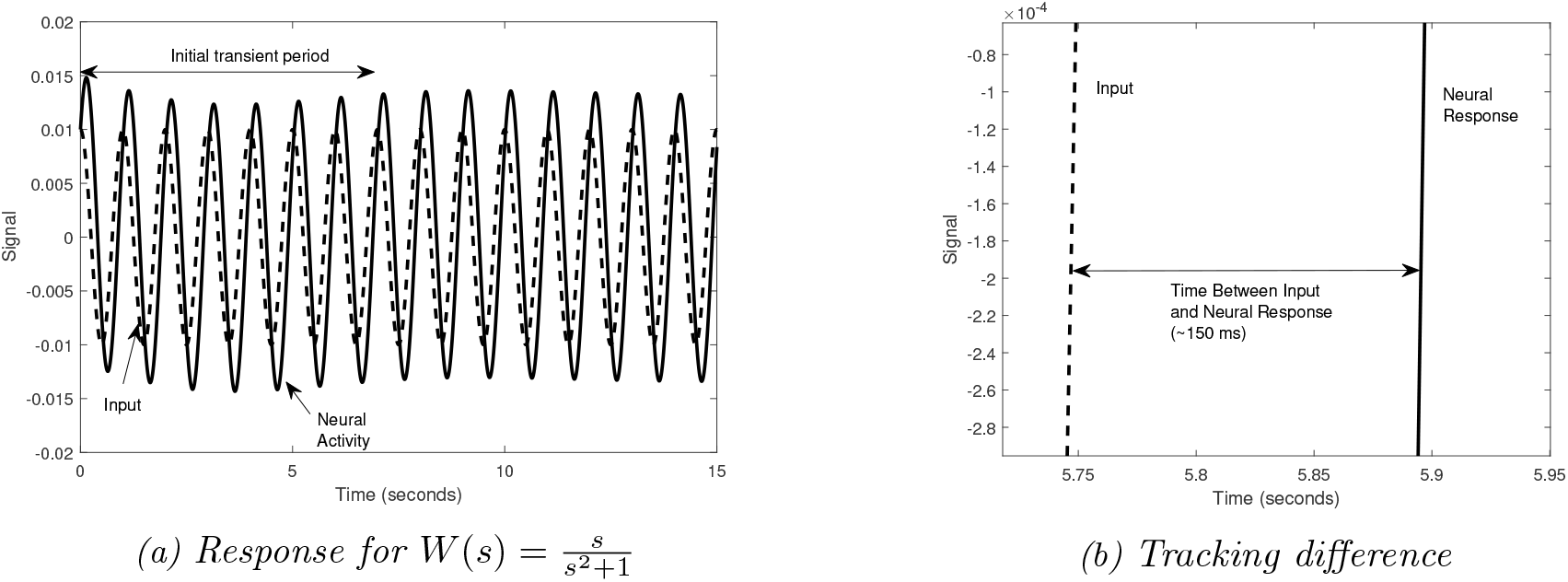
The neural response with an input forcing function that is oscillatory, instead of as an impulse. After an initial transient period, a steady state oscillatory response is seen in which there is a delay between the input signal and the neural signal of approximately 150 milliseconds for this model.

In Figure 8, the response of the neural activity is shown with a forcing function of a sinusoidal wave at 1 Hz. After an initial transient period, there is a steady-state response in which the input signal leads the neuaral response. The amount of lead in time is determined from Figure 8(8a) as approximately 150 milliseconds. This amount is consistent with the time it would take for the input signal to be received, processed through the synaptic cleft, and projected forward one or two connections.

### 3.2 Nonlinear Neurophysiological Activity

A model of nonlinear behavior of fMRI-BOLD signals can characterize the physiological limitations of neuroanatomical features, such as the capability of synaptic cleft and post-synaptic membrane to process neurotransmitters rapidly, and thus such nonlinearities can be used to elucidate the functionality.

In terms of the evidence of nonlinearities of fMRI-BOLD signals, an excellent set of data was presented by Glover [10], who measured the BOLD response in sensorimoter cortices in which the participants tapped fingers while listening to tones of metronome pacing. Glover developed a convolution model data associated with 1 second of effector input, and showed that the BOLD signal did not scale linearly as the effector input increased, and was in fact significantly nonlinear in which Glover noted that a “saturation” effect seemed to be responsible. Glover then went onto to develop a nonlinear modification involving a trapezoidal weighting function to Buxton’s balloon model so as to fit the data.

In this section, the data of [10] is revisited to compare the implementation of the model developed in Section 2 with the previous models, including the popular Balloon model and its nonlinear implementation. Moreover, because of the separation involved with the neural signal *X*(s) and the chemical signal Q(s), an evaluation is conducted of the neural response to handle nonlinear responses. The model describing the data of [10] is given as:

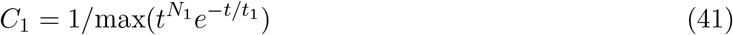

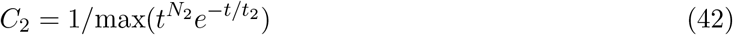

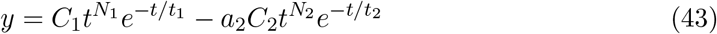

where *N*_1_ = 5, *N*_2_ = 12, *a*_2_ = 0.4, *t*_1_ = 1.1, *t*_2_ = 0.9.

To start the analysis, the linear impulse response function using a 4 pole/4 zero characterization is estimated from the convolution model of Equation 43. In Figure 9, a comparison of the newly derived model to that of the convolution model fit of Equation 43 is presented after an appropriate scaling, indicating excellent agreement at 95.4%.

**Figure 9:**
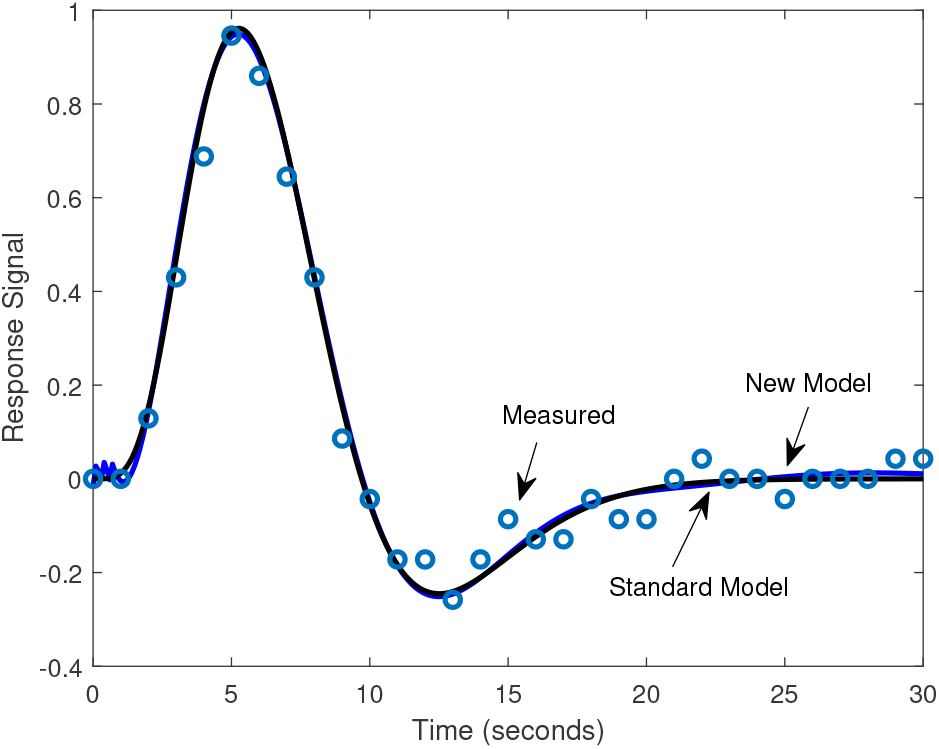
The one second response data from the paper of [10] is compared in relation to the newly developed model. An agreement is shown with a data fit of 95%, and tracks the peak, undershoot and stabilization periods; thus providing evidence of the suitability of the newly developed model.

The resultant model of 4 zeros and 4 poles can then be separated into models of 2 zeros and 2 poles each for *X*(*s*)/*W*(*s*) and the remainder for *Q*(*s*)/*X*(*s*). After a review of the model resulted in conjunction with Equation 26), the representation is as follows,

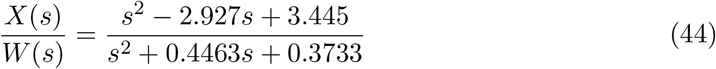

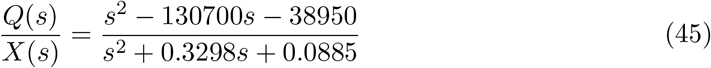

The set of data available from [10] also includes the BOLD measurement after 2, 4, 8 and 16 seconds of finger tapping at 3 Hz, measured over a 30 second interval. If the response was linear, then it would be expected that the resultant convolution would add appropriately, resulting in peaks roughly twice as large for 2 second, in comparison to 1 second, four times as large for 4 seconds, and then 8 and 16 seconds would be five times as large. It would not be more than that because not all inputs add in phase at the later stages of inputs. However, the measured results are much different as will be seen shortly, in which the total range of signals to 16 seconds of input only vary by a factor of approximately 2.

To characterize the nonlinearity according to the methods described in Section 2, a saturation was placed on the scaled neural response of 0.1. In Figure 10, the neural signals associated with the system for 2, 4, 8, and 16 are presented with this cut-off.

**Figure 10:**
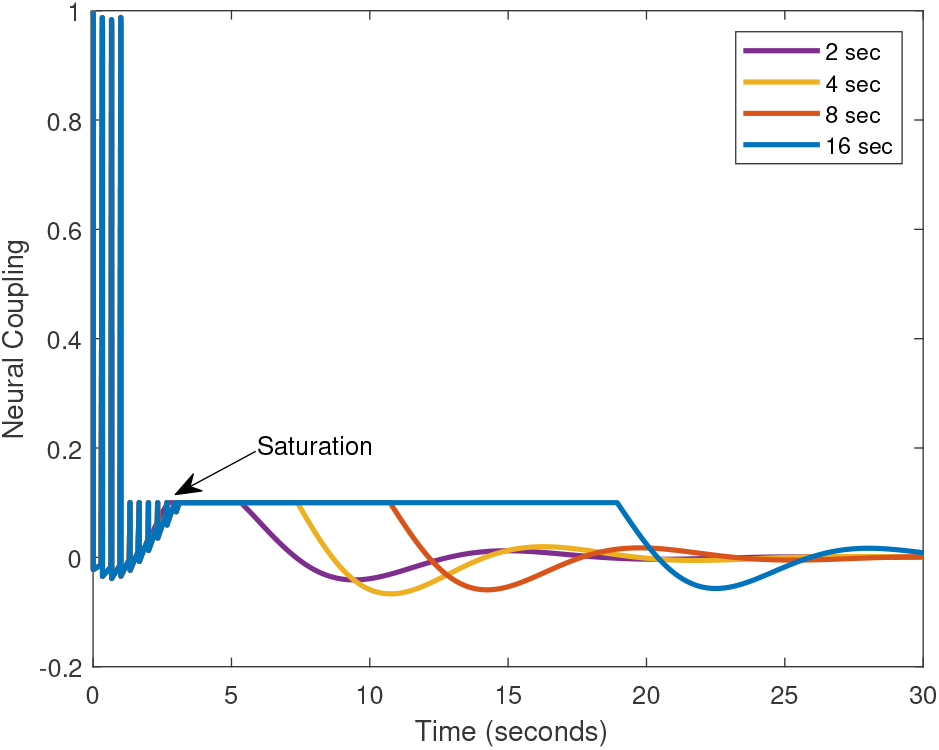
To characterize the nonlinear response of the fMRI signal data set from Glover [10] which shows nonlinearities in amplitude at 2, 4, 8, and 16 seconds, in comparison to one second, the neural signal X(s) of the newly developed model is clipped after 1 second and set to a maximum level of 0.1 for the duration of the experiment. The input signal W(s), a series of impulses at 3 Hz, is run through the model Equation (44) to determine the neural signal, which is then limited indicating saturation of post-synaptic neuron, and the metabolic signal is then run via Equation (45).

After saturation, the decay of the signals are indicated by modeling the trailing edge of the settling period and then added to the signals that start from an unsaturated state according to Equation (35). In Table 1, the models for the decay of the saturated signal are presented.

**Table 1:** The model representations of the response relating to neural and metabolic functions for operation in a saturated membrane regime. Each of the impulse sequences of 2, 4, 8 and 16 seconds are identified.

| Model | 2 | 4 | 8 | 16 |
| --- | --- | --- | --- | --- |
| $\frac{Q(s)}{W(s)}\vert _{\text{sat}}$ | $\frac{-0.44(s-2.1)e^{-2s}}{s^2+0.37s+0.16}$ | $\frac{0.039(s+27)e^{-2s}}{s^2+0.49s+0.15}$ | $\frac{0.46(s+2.8)e^{-2s}}{s^2+0.54s+0.13}$ | $\frac{0.35(s+15)e^{-2s}}{s^2+0.45s+0.085}$ |

To determine the additive function of the saturated and unsaturated signal origin according to Equation (35), a crossing point of 14 seconds was selected and a 90% response time of approximately 10 seconds was used to characterize the sigmoid curve of Figure 11.

**Figure 11:**
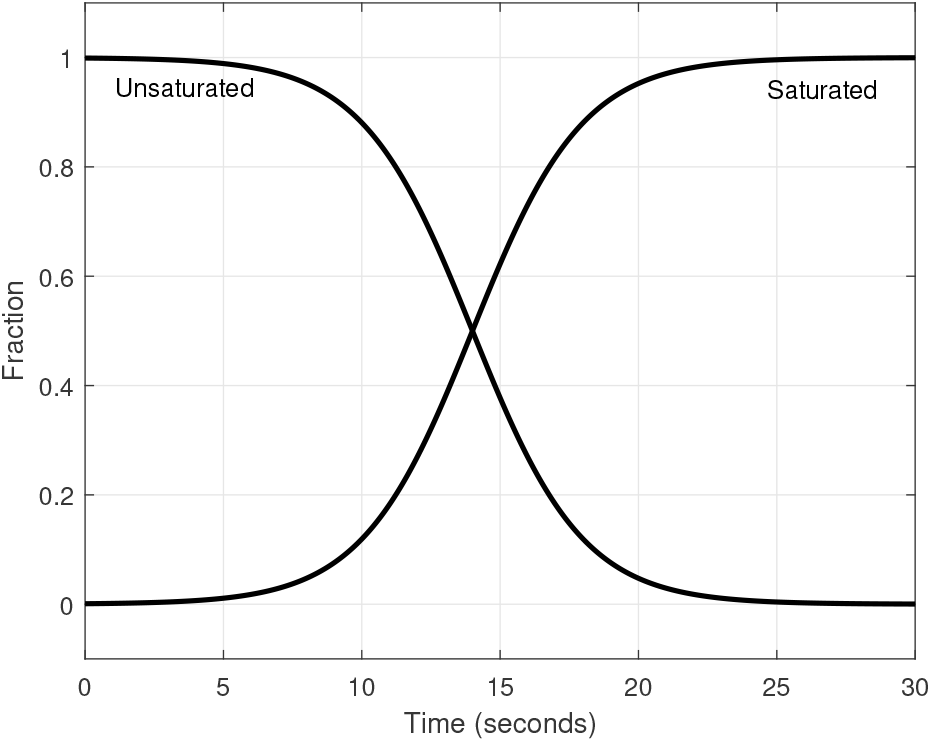
A sigmoid shaped function is used to determine the fraction of signals which begin unsaturated in relation those that emanate from saturated signals. The shape of the function is based off of the facilitated diffusion equation for saturation of the plasma membrane substrate.

After implementation of Equations (28), (37), and (35), the result for the 2, 4, 8 and 16 seconds of pulsed finger tapping is shown in Figure 12 in comparison to the measured data estimated from that of Glover [10].

**Figure 12:**
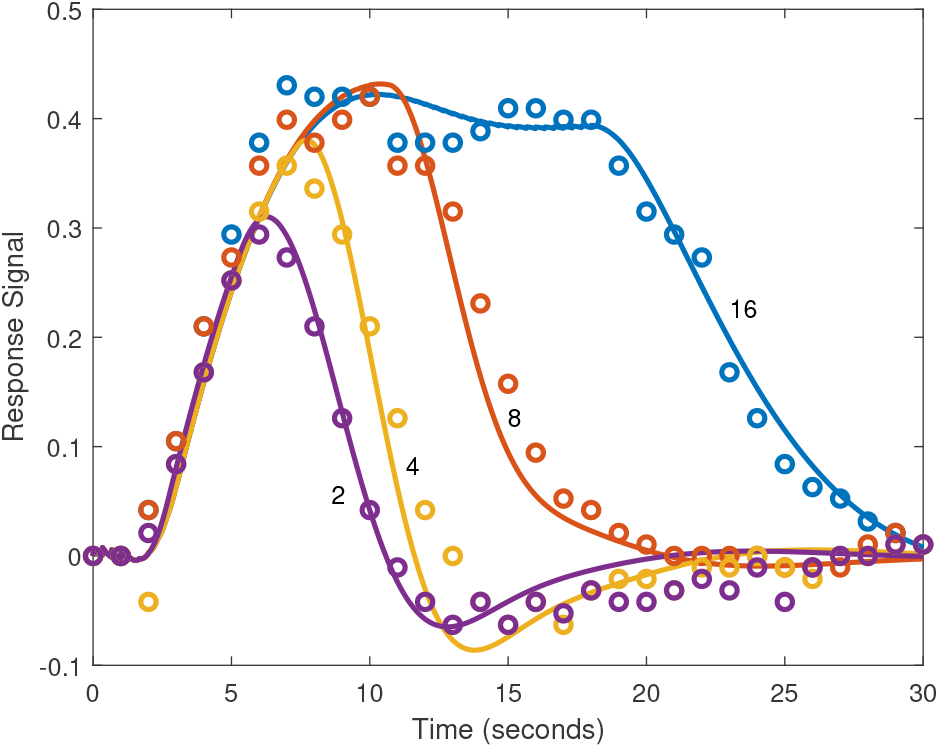
Utilizing the saturated signal of Figure 10 and the percentage of saturation of Figure 11, along with the ramp-down model of Equation (28), is compared against the data of Glover [10] for the nonlinear operational regime of input signal duration 2, 4, 8 and 16 seconds. Note that the model captures the upward amplitude nonlinearity, which is primarily due to the maximum allowable neural signal, and the down slope to stabilization is captured.

The composite of the two functions when the signal begins from an unsaturated to a saturated state is shown in Figure 13 as well as the composite result. The signal for 2 seconds is seen to be almost entirely comprised of the initiating response, which is also the case for 4 seconds of input. The transition for 8 seconds of impulsed input shows a contribution towards the end of the transient as comprised of a saturated and unsaturated inputs. For an input sequence lasting 16 seconds, there is a balanced contribution from the initiating and concluding response curves.

**Figure 13:**
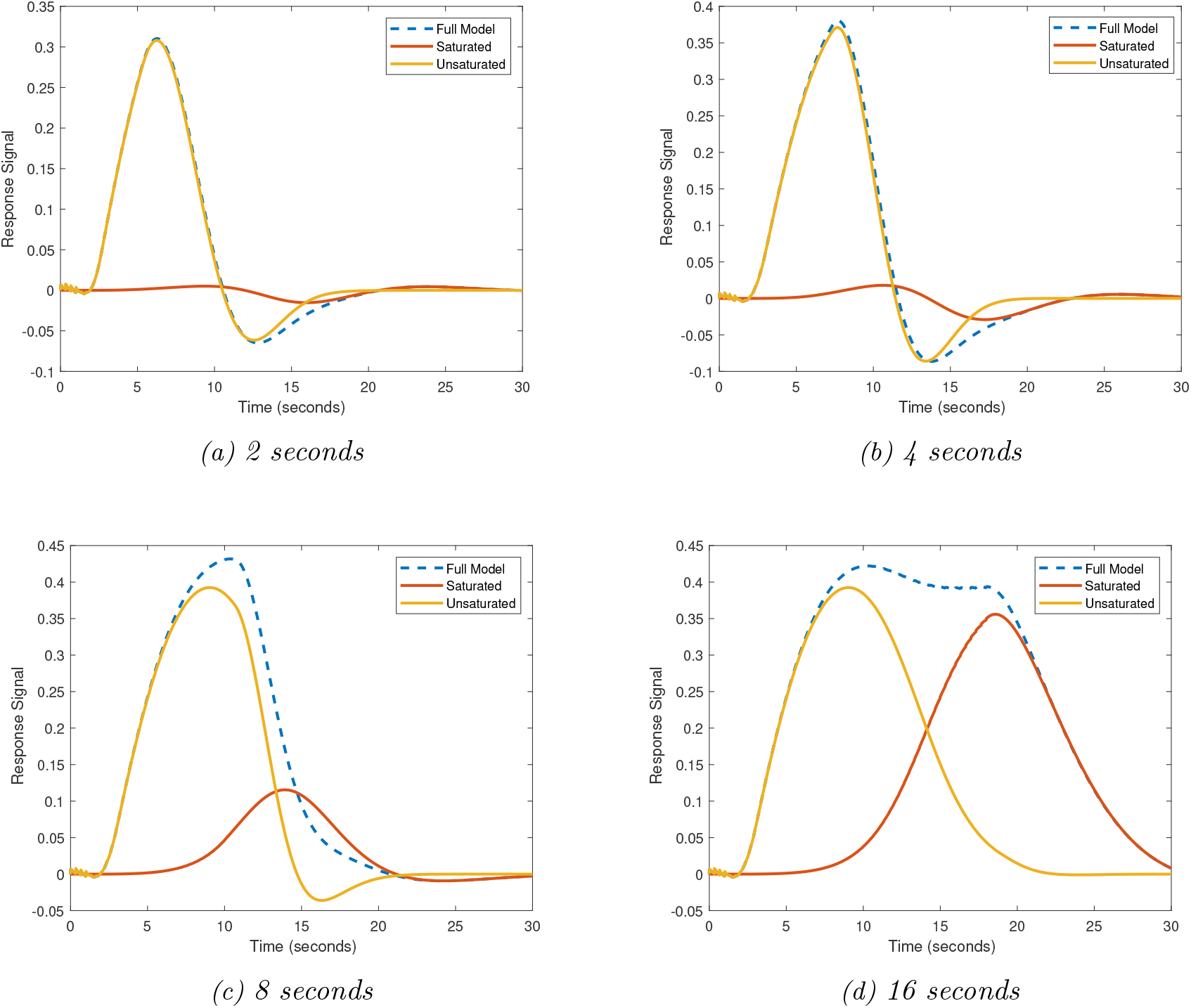
Comparison of the signals which are initiated from an unsaturated and saturated conditions, whose superposition determine the overall BOLD signal response as represented by the dashed curve. Note that the saturated condition explains the downslope in returning to steady-state and the upward rise nonlinearity is explained by the unsaturated initiation, which is ultimately clipped after one second. The nuances of the fMRI signal in between the initial and final steadying can be explained as the integration of the responses as due to the saturation of the neuron, and loss of excitation capability.

The bode plots of several transfer functions are presented in Figure 14 for the relationship of the neural relation *X*(*s*)/*W*(*s*), and the metabolic responses *Q(s)/X(s)* and *Q(s)/W(s)*, along with the transfer function associated with the saturated reduction of *Q(s)* corresponding to an input sequence lasting 16 seconds. The resonance peak of *X(s)/W(s)* is noted relative to the smaller peak of *Q(s)/X(s)*, as well as the absence of the slow frequency contributions, as it is largely comprised of fast frequencies which cuts off corresponding to a time constant of 1.7 seconds.

**Figure 14:**
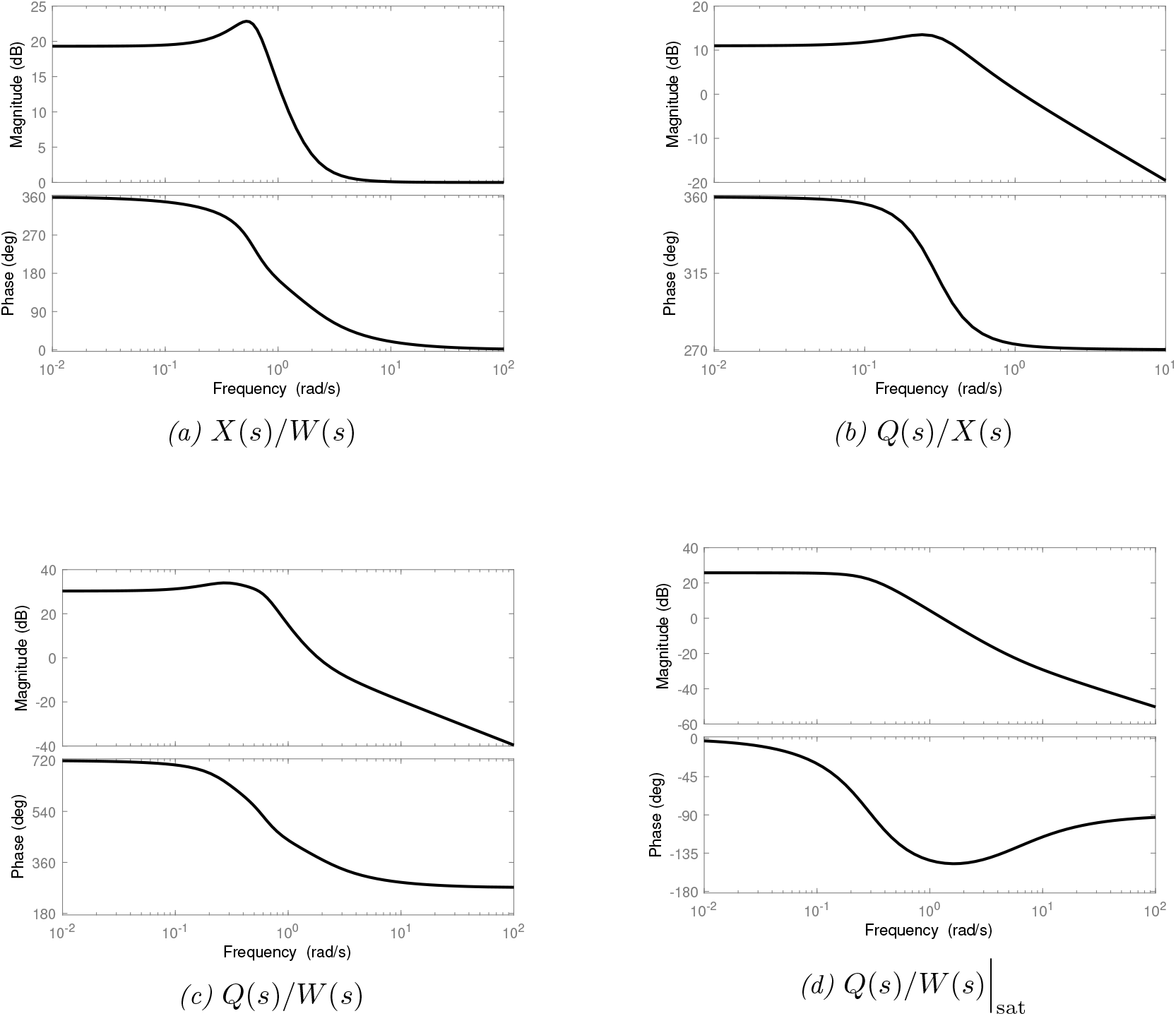
The frequency response of the models characterizing the response for the metabolic and neural signals for the nonlinear cases, as well as the response for the saturated condition during termination for 16 second impulse response. It is noted that there is a strong resonant frequency for the neural signal and a loss of resonant frequency during saturated operation.

## 4 Discussion

In assessing the capability of the model of fMRI BOLD signals, it first is indicated that the model development was derived within a recent framework deployed for examination of neurovascular coupling [29]. In this strategy, the metabolic and neural components are coordinated at the synaptic cleft within the central nervous system, for example at the hypothalamus in the case of body temperature dynamics. The fact that the same framework, which was based on neural signals generated from a combination of neurological and chemical inputs and its rate of change, is confirmatory for both the implementation of fMRI BOLD signals and the framework itself. Of course, additional studies are required to affirm its use, but the predictive capability using this framework appears significant.

As a firm indication of the utility of the modeling approach, the fit of the proposed model to existing models that characterize BOLD signals response is outstanding, as exemplified by the fit of 97.4% to the three gamma model, which has been confirmed itself through comparison to experimental data. Further the parameterization of the model as four zeros and four poles seems appropriate in relation to the underlying theory. In evaluating other model structures, a model with three poles and two zeros indicates a fit of 88.7% to the three gamma model, and a model with five poles and four zeros produces a fit of 99.2 %, which is similar to the fit of four poles, three zeros. It is noted that the use of three zeros versus four zeros produces a negligible loss in accuracy. To further evaluate the capability of the models, the input excitation could be adjusted in amplitude and frequency to improve the model accuracy and predictive capability by enriching the data used for system identification. However, the model captures the important characteristics of the BOLD response including the initial dip, overshoot, undershoot, and stabilization profile.

An important result of this research is the separation of neural activity and metabolic reactivity from the BOLD signals, which is a composite of both sources. The results indicate that the response of neural activity of fMRI signals can be described by a second order plus two zero model in response to a event-driven stimuli. Further, the result indicates that this response is affected not only by the amplitude of the incoming signal changes, but also by its rate of change, as indicated by the Equation (39), which depends upon the proportional and derivative terms of the control law expression for U(s). Upon examination of the neural response model Equation (39) it is noted that the rapid response to the incoming signal is because it is a proper, but not strictly proper, transfer function as the order the numerator is equal to the order of the denominator. This property gives the neural response effectively a derivative action, which makes the actions particularly fast and sensitive to incoming signals, which is desirable for processing stimuli, both internal and external.

In addition, in regard to the model order, it is noted that for the expression for *Q(s)/X(s)* the zero at 4.008 × 10^4^ does not significantly impact the response, and thus the effective model order is 2 poles and 1 zero at least for the impulse response. This makes the response strictly proper for the effect of *Q(s)* on *X(s)*, essentially a responsive reaction to changes in *X(s)*. Thus the neural response would be fueled by metabolic functions which were available at the time of the activity, rather than immediately commandeered. This can further be seen as the response for *Q(s)/X(s)* is a non-minimum phase, or inverse response, and thus the response would be in a consumption mode, followed by reestablishment of the metabolic function. Moreover, both applications studied in the work, the three gamma model and the nonlinear approach, exhibited similar characteristics in *Q(s)/X(s)* with a single zero representation and an inverse response, which aids in confirming the results.

The timescales of the responses are also of interest. The BOLD signal is of course sluggish in that it takes up to thirty seconds to fully settle in response to a signal that may take only a second to excite the system. The action potential of a neural signal is of course very rapid, occuring on the order of 10 to 100 milliseconds; however, for the large input sequences used for fMRI studies in order to generate a significant signal for measurement will necessarily take longer to process since the synaptic processes are a controlling feature. It was noted however that at steady state where the input signals are very small, the model simulations indicate a lag between input and output of roughly 100 milliseconds, which is consistent with measurements of the time scaling needed to generate an action potential.

As the model is an accurate representation of the data for single variable, it is extendable to model sequences of neural actions for multivariable applications. In indicating the utility of the model, it is of interest to note that the neural response has resonant modes, and thus can be sufficiently amplified if the incoming signal is of a similar frequency. This impacts the connectivity of the neural actions and limits the responses which need to be sufficiently damped in order to achieve stable performance. The model indicates that there are only three or four levels possible of connectivity possible if there is a one-to-one connection of neurons. However, to circumvent this issue, the neurons can receive many inputs from sources in order not to excite the resonant modes, but to increase the overall system response. The neural representation developed in this work can be utilized conveniently to study the connectivity since it incorporates linear transfer function models.

The proposed model was extended to nonlinear responses, comparing against alternative models and experimental data, of which the output signals were not simply a linear superposition of input signals. It was necessary to clip the input signal as a saturated function to explain the loss in amplitude while the input stimuli frequency is increased; however, this saturation was based upon the capabilities of the dynamics of the synaptic cleft and receptor saturation as reflected in the neural transfer function. By analysis of the trailing edge of how the neuron response when saturated, the neural signal was predicted as a strictly proper transfer function of two poles and one zero, and thus there is a loss of excitability in comparison to the proper transfer function of two pole, two zero representation for an unsaturated signal.

Furthermore, in comparing the linear and nonlinear responses, the nonlinear responses contain more nuanced responses than the linear ones, which mainly have a dip, overshoot, undershoot, and settling. The nonlinear response retains these features, but also include subtle variations during extended periods of hold and ramp-down, for example a quarter-cycle oscillation as seen for the response corresponding to the 16 second input function or the ramp-down during the 8 second input function. From the characterization, this slow variation is due to the combination of the strictly proper transfer function response of the neural signal in unsaturated mode in relation to the proper transfer function of the saturated signal response. This was modeled by the sigmoid function, which was produced in analysis of the membrane dynamics.

## 5 Summary of Main Results

The main findings of the work are as follows:

- The BOLD signal is accurately modeled by a transfer function comprised of four poles and four zeros for external stimuli triggering a linear response;
- An analytic representation of neural activity can be extracted from a fMRI BOLD as a dynamic transfer function comprised of two poles and two zeros;
- The BOLD signal can be characterized as neural activity, which has faster dyanamics when triggered by external stimuli, driving a metabolic function;
- Because of resonant modes characterizing the neural response, the number of connections to like neurons may be limited;
- The linear model can be extended to nonlinear response conditions by incorporating saturation functions applied to the neural response;
- A framework for modeling neurovascular coupling was demonstrated for fMRI BOLD signals, and such framework was previously demonstrated for autonomic neurovascular coupling within the hypothalamus to model body temperature in response to infectious agents, thus providing additional evidence in support of the framework;
- The salient features of the BOLD signal are captured within the neural response model, such as initial dip, peak response, undershoot, and stabilization and thus it is a conclusoin that neural activity is well characterized by BOLD signals.

## 6 Conclusions

A model has been developed which allows for separable neural and metabolic metrics as transfer functions to describe fMRI BOLD signals. The accuracy of the model to describe linear fMRI signals is excellent in comparing the signal response to the three gamma model, as to ones derived from experimental data. Further, the utility of the model is of a general form that can easily integrate linear time-invariant model analysis, which can be extended to multivariable analysis investigating anatomical coupling of brain function. A control law that included time differentials of chemical and neural signals was incorporated within the model, which yielded a neural signal model that is immediately responsive to incoming stimuli. The extension of the technique to handle nonlinear fMRI BOLD signals was performed, and the detailed features of such nonlinear responses were effectively modeled and noted to be due to the transitions between saturated and unsaturated conditions. A general framework for neurovascular coupling was used in the development of the neural and metabolic models, which thereby assists in the parallel validation of the framework itself. The model description of the neural activity was very similar in shape to the BOLD signal, and thus the conclusion is that the fMRI BOLD signal quantifies the essential components of neural activity.

